# Chirp Sensitivity and Vowel Coding in the Inferior Colliculus

**DOI:** 10.1101/2025.03.10.642479

**Authors:** Paul W. Mitchell, Laurel H. Carney

## Abstract

The inferior colliculus (IC) is an important brain region to understand neural encoding of complex sounds due to its diverse sound-feature sensitivities, including features that are affected by peripheral nonlinearities. Recent physiological studies in rabbit IC demonstrate that IC neurons are sensitive to chirp direction and velocity. Fast spectrotemporal changes, known as chirps, are contained within pitch-periods of natural vowels. Here, we use a combination of physiological and modeling strategies to assess the impact of chirp-sensitivity on vowel coding. Neural responses to vowel stimuli were recorded and vowel-token identification was evaluated based on average-rate and spike-timing metrics. Response timing was found to result in higher identification accuracy than rate. Additionally, rate bias towards low-velocity chirps, independent of chirp direction, was shown to correlate with higher vowel-identification accuracy based on timing. Also, direction bias in response to chirps of high velocity was shown to correlate with vowel-identification accuracy based on both rate and timing. Responses to natural-vowel tokens of individual neurons were simulated using an IC model with controllable chirp sensitivity. Responses of upward-biased, downward-biased, and non-selective model neurons were generated. Manipulating chirp sensitivity influenced response profiles across natural vowel tokens and vowel discrimination based on model-neuron responses. More work is needed to match all features of model responses to those of physiological recordings.

**Highlights:**

- Due to phase differences between harmonics caused by vocal-tract resonances, spoken vowels contain fast (within-pitch-period) frequency sweeps, or chirps, to which neurons in the inferior colliculus (IC) are sensitive.
- Both vowel responses and chirp-velocity sensitivity were recorded from single neurons in the IC.
- An analysis of responses from a large number of IC neurons showed that features of chirp sensitivity were correlated to the accuracy of vowel identification based on average rate and/or timing of neural responses.
- An IC model with chirp sensitivity was used to explore its impact on vowel responses.

## 1. Introduction

Vowels carry vital linguistic information and are thus an important signal for speech perception (Kewley-Port et al., 2007). Acoustically, vowels are defined by the shape of a harmonic spectrum and by the fundamental frequency, F0, which is determined by the pitch of the speaker’s voice. Vowels can be distinguished from one another by the frequencies of the spectral peaks, or formants, resulting from vocal-tract filtering (Fant, 1960). The first two formant frequencies, referred to as F1 and F2, are sufficient for vowel identification (Fant, 1960; Hillenbrand et al, 1995). It is important to understand how the auditory system converts these acoustic spectral peaks into a neural code by which vowel discrimination is accomplished. However, non-linearity in the auditory system shapes neural representation of these signals such that the underlying mechanisms supporting vowel coding at various levels of the auditory system are not well understood. Traditionally, studies of speech coding in the auditory nerve (AN) have established average rate and temporal fine structure to be primary candidates for vowel identification (Sachs and Young, 1979; Young and Sachs, 1979), but both of these codes fail at moderately high sound levels (Delgutte and Kiang, 1984a) or in background noise (Delgutte and Kiang, 1984b), suggesting that a more comprehensive vowel code may take shape elsewhere in the auditory system.

The inferior colliculus (IC) is a nearly obligatory synapse in the ascending auditory pathway (Aitkin and Phillips, 1983). In addition to spectral tuning to a characteristic frequency (CF), IC neurons commonly display sensitivity to amplitude modulation (AM) (Schreiner and Langner, 1997; Joris et al., 2004). Modulation transfer functions (MTFs), which depict neural rate versus modulation frequency (Joris et al., 2004), reveal that the best modulation frequencies (BMFs) of IC neurons span the F0-range of speech (Langner, 1992; Krishna and Semple, 2000; Kim et al., 2020). Sensitivity to F0 periodicity has been suggested to underlie a representation of vowels in the IC by converting neural fluctuations in the periphery to rate coding of formants (Carney et al., 2015; Carney, 2018, 2024), and is an example of how speech coding can emerge from sensitivity to sound features.

Beyond AM tuning, most IC neurons are also sensitive to the velocity of fast frequency chirps resulting from phase-differences between components in harmonic sounds (Steenken et al., 2023; Henry et al., 2023; Mitchell et al., 2023). This sensitivity is reflected in rate differences in response to chirps having opposite directions but identical speeds. Sensitivity to chirp direction has been observed in the majority of IC neurons tested (Mitchell et al., 2023). The implications of chirp-velocity sensitivity for vowel processing are unknown. Phase-differences between harmonics, which are associated with frequency chirps, are a feature of natural speech, due to the resonant characteristics of the vocal-tract filter. A related feature of vowels, the group-delay functions that track time delay versus frequency, indicate peaks in time lag of harmonics at and around formant frequencies (Bozkurt et al., 2006; Rajan et al., 2013).

In this paper, we test the hypothesis that IC chirp sensitivity is related to neural coding of vowels using physiological and modeling methods. Examining the responses of IC neurons to natural vowels, we evaluate the correlation of chirp-direction bias and average rate and temporal coding of vowels. Finally, using a physiologically plausible model of chirp-sensitive IC neurons, we simulate and compare responses to vowel stimuli of model cells with and without chirp sensitivity.

## 2. Methods

To test the hypothesis that chirp-sensitivity of IC neurons is correlated to vowel responses, we employed a combination of physiological and modeling methods.

### 2.1 Neural Recordings

Physiological methods are described in detail in Mitchell et al. (2023). Briefly, extracellular neural recordings were made in the central nucleus of the inferior colliculus (ICC) of awake Dutch-belted rabbits (Oryctolagus cuniculus) using tetrodes. Data were collected in a total of 5 animals with normal hearing, assessed using distortion product otoacoustic emissions (DPOAEs). All methods were approved by the University of Rochester Committee on Animal Resources.

During an initial surgery, a custom 3D-printed plastic headbar was attached to the skull. After healing, a craniotomy (∼2 mm diameter) was made to access the IC and a microdrive (Five-drive, Neuralynx, Inc., Bozeman, MT, USA) was mounted on the headbar to advance implanted tetrodes through the ICC. Tetrodes, each consisting of four 18-μm platinum-iridium epoxy-coated wires (California Fine Wire Co., Grover Beach, CA, USA) and plated with platinum black to obtain impedances of approximately 0.1 – 0.5 MOhms, were replaced every 1-3 months during additional surgical procedures. All surgeries were performed under anesthesia (ketamine (66 mg/kg) and xylazine (2 mg/kg), administered intramuscularly). Data collection was conducted daily in 2-hour sessions in a sound-attenuated booth (Acoustic Systems, Austin, TX, USA). Sound was delivered to the rabbits via custom-made earmolds Custom earmolds (Dreve Otoform Ak, Unna, Germany). At the beginning of every session, the acoustic system was calibrated (ER-7C probe-tube microphone, Etymotic Research, Inc., Elk Grove Village, IL) to compensate the stimuli for the frequency response of the system.

Voltage recordings were made using a multi-channel recording system (RHD system, Intan Technologies, LLC., Los Angeles, CA, USA) and Intan software. To identify single-unit action potentials (spikes), the voltage recording was filtered using a 4th-order Butterworth bandpass filter (300 – 3000 Hz). Spikes were identified when the voltage recording exceed a threshold defined as four standard deviations of the signal. Features of the spike waveforms were used to sort spikes into clusters, primarily the slope of repolarization (Schwarz et al., 2012). Clusters were identified as a single-unit neural response when less than 2% of the inter-spike intervals were shorter than 1 ms. Only single-unit recordings were included in the analyses presented below. Neurons identified in consecutive sessions were considered unique only if both tetrode location and response properties changed.

### 2.2 Stimuli

Frequency response maps (RMs) were used to assess CF, the frequency that elicited the highest response rate near threshold. To generate RMs, a series of 0.2-s-duration tones were presented at different levels (13-, 33-, 53-, and 73-dB SPL) and frequencies (250 Hz—16 kHz) in random order. Each tone was presented 3 times, either contralaterally or diotically, and included 10-ms raised-cosine on/off ramps. Tones were separated by 0.4 s of silence.

MTFs were used to assess neural sensitivity to 100% sinusoidally AM wideband noise (100 Hz - 10 kHz; 1-s duration including 50-ms on/off cos^2^ ramps, presented diotically at 33-dB SPL spectrum level, overall level of 73 dB SPL). Modulation frequencies spanned from 2-350 Hz, with 3 steps per octave. Each modulation frequency was presented a total of 5 times in random order, and stimuli were generated for each trial. MTFs were categorized based on shape. The results presented below focus on examples from the two largest groups of MTF types: band-enhanced (BE) neurons, which have significantly increased rate for a band of modulation frequencies relative to unmodulated rate, and band-suppressed (BS) neurons, which have significantly decreased rate for a band of modulation frequencies relative to unmodulated rate (Kim et al, 2020; Mitchell et al., 2023).

Rate-velocity functions (RVFs) were used to evaluate chirp-sensitivity of IC neurons (Mitchell et al., 2023). A set of chirps was generated with velocities ±0.40, ±0.80, ±1.59, ±3.16, ±6.24, and ±9.24 kHz/ms (equivalent to Schroeder-harmonic complexes with F0s equal to 25, 50, 100, 200, 400, and 600 Hz, respectively, and with instantaneous frequencies spanning from F0 to 16 kHz). To normalize energy in these variable-duration chirps, stimuli were assigned a sound level equal to 68 dB SPL - 10×log_10_(*T*/*T*_ref_), where *T* is the duration of the chirp, and *T*_ref_ = 2.5 ms (duration of the chirp of ±6.24 kHz/ms). Additionally, cos^2^ on/off ramps were applied with duration equivalent to 10% of chirp duration. Chirps were presented in random order, separated by random intervals of silence ranging from 40 - 60 ms to ensure aperiodicity. Each chirp was presented a total of 840 times to collect one RVF. Response rate was calculated by summing spikes over a 15-ms time window starting at an estimate of the neural latency based on the response to a 73-dB SPL tone at CF (from the response map).

Vowel stimuli were from Hillenbrand et al. (1995), a database that contains recordings of spoken English vowels with a range of average F0s. All twelve vowels in the database were presented, including /iy, ih, ei, eh, ae, ah, aw, oo, uw, er, oa, uh/. The 200-ms steady-state center portion of each vowel was extracted and 25-ms cos^2^ on/off ramps were applied. Stimuli were presented diotically, at 68 dB SPL, in random order, with 30 repetitions per vowel. Responses presented here were for three speakers (identified as “M03”, “M40”, and “W39” in the Hillenbrand dataset), who had average fundamental frequencies of 95, 148, and 202 Hz, respectively.

### 2.3 Vowel-Component Decomposition

To evaluate the chirp cues contained in spoken vowels, the magnitude and phase spectra of the stimuli were estimated, as follows (Yasi, 2004; Ramamurthy and Raghavan, 2013): First, each 200-ms-duration vowel stimulus was divided into four 50-ms segments, and the Welch power spectrum was evaluated for each. The peaks of the power spectrum were identified up to 3.5 kHz; these peak frequencies were the initial estimates of harmonic components. Note that to reject spurious, non-harmonic peaks, power-spectrum peaks that differed by less than 65% from the average F0 were removed. Next, a set of bandpass filters was designed to isolate each harmonic. The Matlab function “kaiserord” was used to generate Kaiser-window filter parameters for a finite impulse response (FIR) filter. The Kaiser window was chosen for its linear phase response. The Matlab function “fir1” was used to generate the filters. Filters were designed with a 20-Hz passband centered around each component frequency, and stopband cutoffs at +/−F0/2 with respect to the component frequency. The response of each filter to the original signal was approximately the waveform of the isolated harmonic component. Using the Matlab function “fmincon”, the frequency, magnitude, and phase of the harmonic were estimated, using the sum of squared errors as the objective function and a starting value of the frequency based on the power spectrum peak. To prevent phase optimization from becoming stuck at the bounds, phase values were constrained between −2π and 2π, and were later assigned equivalent phases between -π and π.

The analysis was repeated for each peak in the power spectrum up to 3.5 kHz to estimate the vowel spectrum. To illustrate the fast frequency chirps within the vowel stimuli due to phase transitions in the spectra near formants, a synthetic version of the vowel was generated that had a uniform magnitude spectrum and the estimated phase spectrum. To illustrate the within-pitch-period frequency chirps in the resulting synthetic vowels, spectrograms were generated using the Matlab function “spectrogram” with a Hamming window over 600-sample segments, allowing for 590 samples of overlap between segments, with an overall sampling rate of 48,828 samples/sec. Finally, formant frequencies of the Hillenbrand vowels were identified using Praat analysis (Boersma and Weenink, 1992-2024).

### 2.4 Physiological Response Analysis and Classification

To assess identification of vowels for a given F0 (speaker) based on IC responses, two confusion matrices were constructed, one based on average rates and one on temporal information. The average rate in response to a given repetition of one vowel was calculated, and the overall average rates across repetitions in response to each of the 12 vowels was calculated, excluding the current repetition. The response to each repetition was identified as the vowel for which the absolute difference between the single-repetition rate and overall average rate was minimal.

Vowel identification based on spike timing used a strategy developed by Satuvuori et al. (2017) called rate-independent spike (RIS) distance, a measure of the temporal similarity of two spike trains that is unaffected by average rate. Briefly, the method involves matching each spike to its temporally closest neighbor in the other spike train. A profile of each spike train is constructed describing temporal similarity to the other train on a sample-by-sample basis. Averaging the two profiles and dividing by the local average rate of the two spike trains yields a final distance estimate (Satuvuori et al., 2017). The RIS distance is a value between 0 and 1 that evaluates similarity solely based on spike timing (Satuvuori and Kreuz, 2018). For vowel identification, the RIS distance for every possible pair of spike trains was calculated. Each repetition was identified with the vowel for which the average RIS distance was smallest, excluding self-comparisons.

For both methods, a confusion matrix was generated by tallying the identification results on true-vowel vs. predicted-vowel axes. Vowel-by-vowel accuracy was assessed using the formula

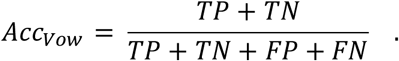

Here, true positives (TP) were the number of correct predictions of the vowel (the square on the diagonal). True negatives (TN) were the number of correct rejections (all other squares on the diagonal). False positives (FP) were the number of incorrect predictions of the vowel (all squares in the vowel’s column except the diagonal). False negatives (FN) were the number of incorrect rejections of the vowel (all squares in the vowel’s row except the diagonal). Additionally, the overall accuracy of identification was calculated by dividing the total number of correct identifications (sum of values in squares on the diagonal of the confusion matrix) by the total number of all identifications (sum of all squares).

Overall identification accuracy based on the physiological responses was then related to the chirp sensitivity of the neurons. To extract prominent features of chirp sensitivity from the population of RVFs, as in Mitchell et al. (2023), principal component analysis (PCA) was conducted using the Matlab function “PCA”. RVFs were normalized by individual peak rates before analysis. Note that principal component analysis performed in this study used a different set of neurons than those used in Mitchell et al. (2023), although there was some overlap between datasets. Thus, the exact shape of the first three principal components is not the same between the two studies. However, the resulting principal components are similar between studies, thus the same interpretation could be applied to the results of the PCA analysis for both sets of neurons.

Additionally, a metric based on receiver-operating characteristic (ROC) analysis (Egan, 1975) was used to measure an index of direction selectivity in high-velocity (> 2 kHz/ms) chirp responses, independent of direction. ROC was used to measure the discriminability of chirp direction based on single-chirp response rates for chirps of equivalent speeds (absolute value of velocity) but opposite directions. Calculating the area-under-the-curve of the resulting ROC function gave the direction bias per velocity pair as a number from 0 to 1, where 0 indicated downward bias, and 1 indicated upward bias. To obtain high-velocity direction bias, the direction bias of the velocities greater than 2 kHz/ms (3.16, 6.24, and 9.24 kHz/ms) were averaged. Then, high-velocity direction bias was calculated as 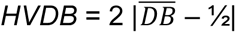, where 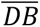 was the mean of direction biases. High-velocity direction bias described direction bias, independent of direction, on a scale from 0 to 1, with 1 indicating strong direction bias.

### 2.5 Modeling Vowel Responses with a Chirp-Sensitive IC Model

The model for IC-cell chirp-sensitivity is detailed in Mitchell and Carney (2024). Briefly, the mechanism of the model depends on a chirp-direction-sensitive inhibition of the IC originating from octopus cells of the posteroventral cochlear nucleus (PVCN), relayed through the ventral nucleus of the lateral lemniscus (VNLL). The model uses a strategy described by Krips and Furst (2009), in which non-homogeneous Poisson processes (NHPPs) represent excitatory or inhibitory inputs to coincidence detectors (CDs), which have responses that are also described by NHPPs (Krips and Furst 2009). The responses of IC neurons were modeled starting from model AN fibers to the model IC neuron, with inhibitory and/or excitatory inputs to CDs at each stage of the model (Mitchell and Carney, 2024).

In the chirp-sensitive model has two stages: an octopus-cell stage and an IC stage. The octopus-cell stage receives AN inputs tuned to two different frequencies, on-CF (the CF of the final IC neuron) and off-CF (OCF). Single AN inputs are subthreshold for the model octopus neuron. By increasing the number of identical CF inputs, N_CF_, the amplitude of the CF input channel is effectively increased to make it suprathreshold. Additionally, all AN inputs to the model octopus neuron elicit a delayed hyperpolarization, implemented as a delayed inhibitory input to the octopus cell. The hyperpolarization suppresses the octopus cell equivalently for both CF and OCF inputs. Thus, a mechanism that detects the sequence of the input times emerges (Lu et al., 2022): a chirp eliciting the suprathreshold CF input first is more likely to elicit a response from the octopus cell, whereas a chirp eliciting the subthreshold OCF input first is less likely to result in a response because the delayed OCF hyperpolarization suppresses the later-arriving CF excitation.

The IC model stage receives one on-CF excitatory input, and two inhibitory inputs: one from the octopus-cell model and one delayed input, relayed from an on-CF AN model. The on-CF excitation and inhibition are delayed by delay parameters d_E and d_I, where the difference between the two tunes the final neuron’s MTF in a manner described by the same-frequency inhibition-excitation (SFIE) model (Nelson and Carney, 2004). The output of the IC-model stage has chirp-sensitivity opposite to that of the inhibitory octopus stage and has a BE MTF (Mitchell and Carney, 2024).

To test the hypothesis that chirp sensitivity impacts vowel identification based on neural responses, six model neurons were created, three with 2-kHz and three with 1-kHz CFs, for comparison to IC recordings with similar CFs. For both CF groups, one neuron had upward-chirp bias, one had downward bias, and one was non-selective. These upward and downward direction biases resulted from different octopus-model parameters (Table I); the non-selective model had no octopus-model input. Note that the upward- and downward-biased models with 1-kHz CF were identical to those in Mitchell and Carney (2024; their Fig. 4). Additionally, model rate profiles across vowels were normalized to match physiological examples: the 2-kHz models were matched to example IC neuron 1 (see below), which had average vowel rate of 127 spk/s; the 1-kHz models were matched to example IC neuron 4, with average vowel rate of 88.50 spk/s). Rate multipliers for each of the six models are reported in Table I. To apply the RIS metric to model neurons, spike times were generated based on a dead-time (1 ms) modified Poisson process that was generated based on the model response, which was provided as a rate function.

**Table I.** Parameter values for model neurons. Non-selective (NS) neurons did not receive inputs from model octopus neurons, thus not-applicable parameters are listed as “N/A”.

| CF (kHz) | Direction bias | OCF (kHz) | $d_{CF}$ (ms) | $d_{OCF}$ (ms) | $N_{CF}$ | $M_{Oct}$ | $M_I$ | $d_E$ (ms) | $d_I$ (ms) | $\Delta_{Oct}$ (ms) | $\Delta_I$ (ms) | Rate Mult. |
| --- | --- | --- | --- | --- | --- | --- | --- | --- | --- | --- | --- | --- |
| 2 | Up | 1.00 | 0 | 0.45 | 3 | 10 | 10 | 1.2 | 3.7 | 1 | 1 | 4.77 |
| 2 | Down | 3.00 | 1.00 | 0 | 3 | 10 | 10 | 1.2 | 3.7 | 1 | 1 | 4.16 |
| 2 | NS | N/A | 0.30 | 0 | N/A | 0 | 10 | 1.2 | 3.7 | 1 | 1 | 2.61 |
| 1 | Up | 0.90 | 0 | 0.45 | 4 | 12 | 16 | 1.4 | 3.4 | 1 | 1 | 4.80 |
| 1 | Down | 2.21 | 1.00 | 0 | 4 | 12 | 8 | 1.4 | 3.5 | 1 | 1 | 2.31 |
| 1 | NS | N/A | N/A | N/A | N/A | 0 | 8 | 1.4 | 3.9 | 1 | 1 | 2.37 |

## 3. Results

To illustrate chirps that occur in spoken vowels, center-tokens of Hillenbrand-vowel stimuli were analyzed using component-phase decomposition (Yasi, 2004; Ramamurthy and Raghavan, 2013). Magnitude spectra (left) and several pitch periods of synthesized vowels (right) produced by this analysis (/aw/, /ih/, and /uw/, all 95-Hz-F0 speaker) show that for each of the formant frequencies, F1-F3, there is an inflection in the vertical white bands of the synthesized vowel, indicating a local chirp (or frequency sweep) (Fig. 1). This inflection is characterized by positive (upward) chirps just below the formant frequencies and/or negative (downward) chirps just above the formant frequencies—this pattern is consistent with peaks in group delays that have been reported to occur near formant frequencies (Bozkurt et al., 2006; Rajan et al., 2013). Note that a synthetic vowel with zero-phase harmonics appears as a series of vertical lines in this visualization, reflecting no chirp features (not shown).

**Figure 1.**
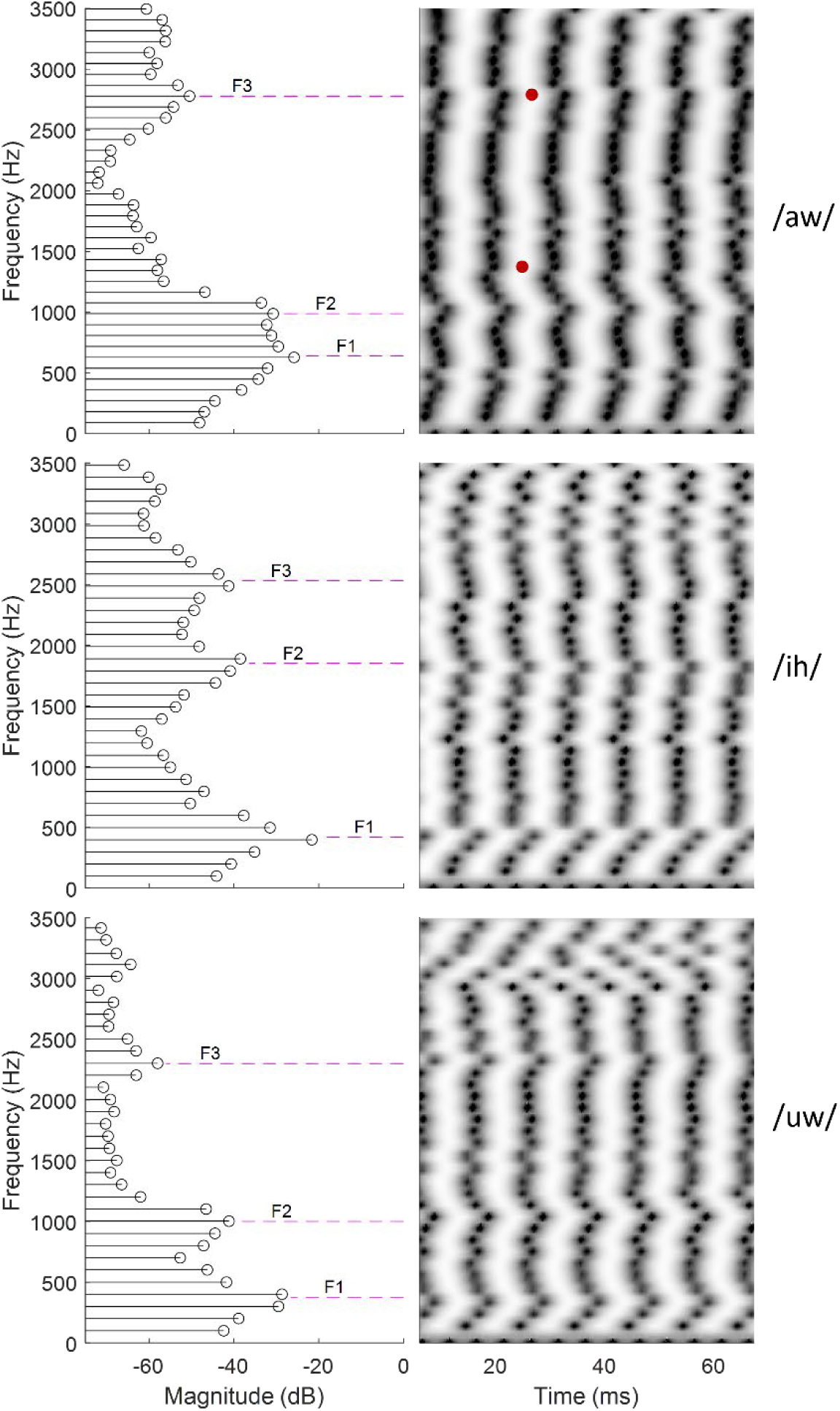
Component-phase decomposition analysis for three representative vowels, /aw/, /ih/ and /uw/, from a speaker with 95-Hz F0. Left) Magnitude spectra of natural vowel tokens resulting from vowel component decomposition. Magenta dashed lines indicate formant frequencies F1, F2, and F3 (matched by frequency to the corresponding synthetic vowel). Right) spectrotemporal patterns in vowels re-synthesized based on results of component decomposition (timing of peaks in each harmonic component are indicated by dark dots). The magnitude spectra of these synthesized vowels were flat. Red dots highlight chirps spanning from 1.38 kHz to 2.81 kHz (see text).

Chirps associated with the inflections near formants are relatively low-velocity and span a limited frequency range as compared to the chirping stimuli used in previous physiological studies, such as Schroeder complexes and aperiodic chirp stimuli (Mitchell et al., 2023); however, the frequency regions between formant inflections include relatively fast chirps that are more comparable to these stimuli. For example, the chirp from 1.38 kHz to 2.81 kHz, between F2 and F3 in the vowel /aw/ (Fig. 1a, red dots) represents a chirp that spans 1.43 kHz over 1.80 ms. The local velocity of this chirp is thus 0.79 kHz/ms, comparable to chirps at the low-velocity end of RVFs used in Mitchell et al. (2023), which ranged from 0.80 to 9.24 kHz/ms. However, note that local velocities vary between F2 and F3, sometimes changing directions and thus approaching near-vertical slopes. Therefore, 0.79 kHz/ms reflects the average chirp velocity between these two frequencies, with local chirp velocities exceeding this average.

Much like the representative vowels in Fig. 1, all vowel stimuli had chirp inflections at formant frequencies, and had faster chirps between formants with velocities equivalent to those used in RVFs. These spectrotemporal features represent a rich set of cues, or spectrotemporal texture, to which chirp-sensitive neurons may be sensitive. Response properties of individual IC neurons to sound features such as frequency, AM, and spectrotemporal chirps are diverse; their responses to vowel stimuli are expected to depend on a combination of these feature sensitivities. Figure 2 shows the response profiles of five example neurons, including CF, MTF, RVF, and vowel rate profiles, alongside vowel confusion matrices based on average rate and timing. These neurons are useful case studies to illustrate how response properties may contribute to vowel identification.

**Figure 2.**
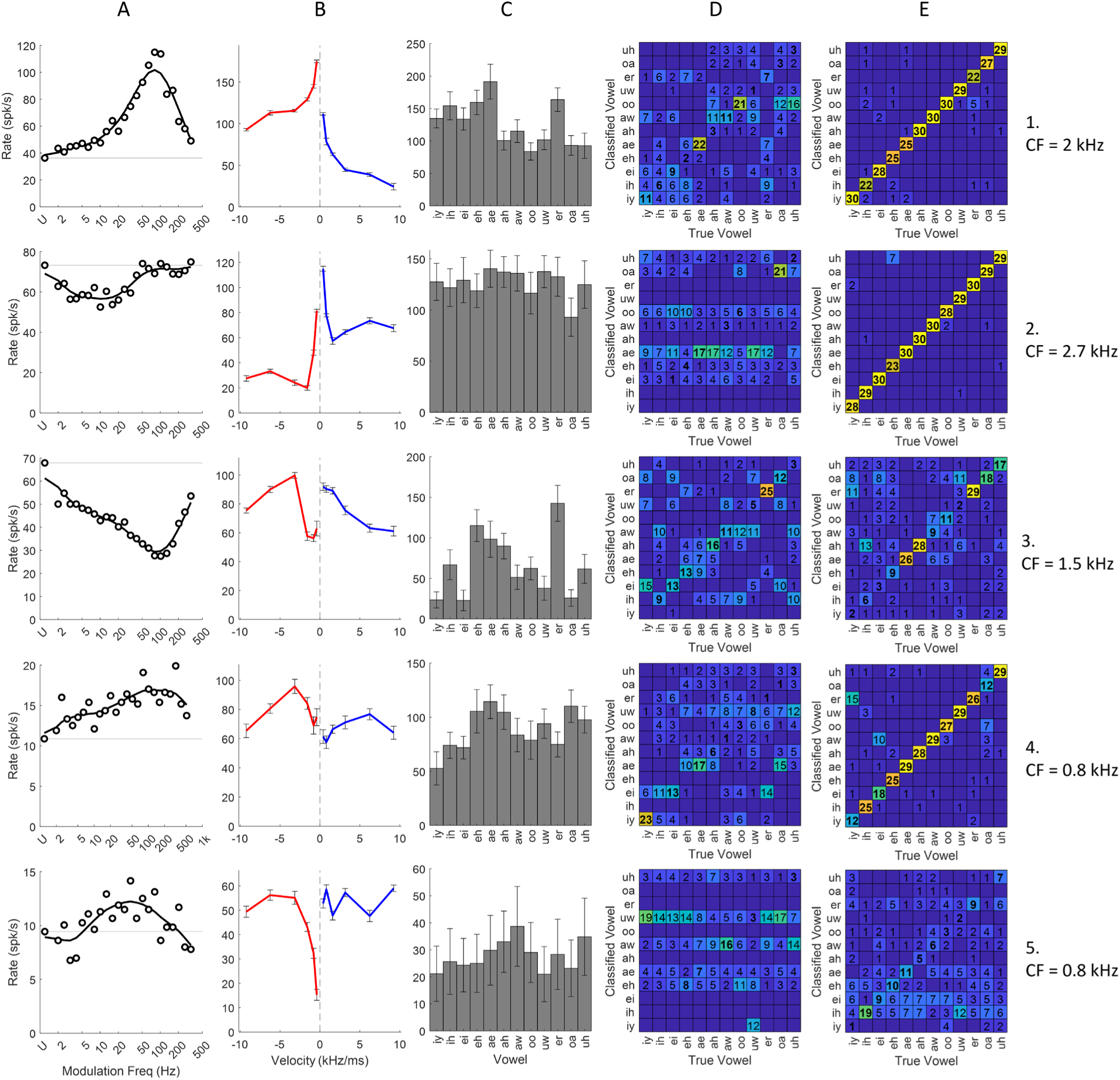
Response profiles of five representative neurons with CFs in the typical F1-F2 range (< 3 kHz). Column A: MTFs based on AM noise. The U on the x-axis indicates unmodulated noise. Solid black curve depicts smoothed MTF. Column B: RVFs, red indicates average-rate responses to downward, and blue to upward chirps; error bars are standard errors. Vertical dashed gray line at 0 velocity. Column C: Average rate histograms in response to 30 repetitions of twelve vowels, 95-Hz F0, from Hillenbrand et al. (1995). Error bars are standard deviations. Columns D and E: Confusion matrices based on average rate (Column D) and timing (Column E) metrics. Numbers of identifications (out of 30 possible) are indicated by boxed numbers; boxes with no numbers were 0. True positive identifications along the diagonal are in bold text.

CFs of the selected neurons were below 3 kHz, in the frequency range of F1 and F2. Example neurons 1, 4 and 5 had BE MTFs, and examples 2 and 3 had BS MTFs (Fig. 2A). Example neuron 1 had an RVF with a consistent and strong bias toward downward chirps, while neuron 2 had a consistently upward-selective RVF (Fig. 2B). Neuron 3 had an RVF with mixed chirp-direction sensitivity, changing direction bias at 2 kHz/ms. Finally, neurons 4 and 5 had identical CFs but opposite RVF direction-bias: neuron 4 was downward biased and neuron 5 was upward biased. These example neurons are representative of the population studied, in that upward or downward chirp-direction bias was not more common among neurons of a specific MTF shape or CF (Mitchell et al., 2023).

Vowel responses are expected to be influenced by proximity of CF to formant frequencies, by the relationship between F0 and peaks or valleys in the MTF, as well as by neural fluctuations related to formant frequencies (e.g., Carney et al., 2015). Lastly, given the presence of chirps similar to those in Fig. 1, an assortment of chirp cues may also influence responses to vowels. To assess vowel discrimination based on IC responses, confusion matrices were generated for sets of vowels with each average F0 and for two neural metrics, average rate and timing. The confusion matrices in Fig. 2 are based on vowels with an average F0 of 95 Hz. Confusion matrices based on average rate (Fig. 2D) show results of vowel identification based on single-repetition responses. Identification performance can be assessed by the true positives (denoted by bold text). Comparison of confusion matrices to the average rate profiles (Fig. 2C) show that the rate classifier did well for vowels that elicited rate extrema. For instance, the response of neuron 3 accurately identified the vowel /er/ due to its high average response rate to that vowel. The response of neuron 4 accurately identified the vowel /iy/ due to its low average response rate to that vowel. Average-rate-based identification was comparatively inaccurate for those vowels that were not extrema.

Vowel identification based on the RIS temporal metric (Satuvuori et al., 2017) generally outperformed that based on average rate, with individual accuracy reaching 100% (all 30 repetitions correctly classified) for many vowels (Fig. 2E). These confusion matrices indicate that for many neurons (examples 1, 2, 4) a vowel-coding scheme based on timing would correctly identify all twelve vowels.

Confusion matrices can be summarized using overall accuracy, which sums the total number of correct identifications (bold text) and divides by the total number of identifications. Using overall accuracy, the performance of timing-based identification of example neurons 1, 2 and 4 (overall accuracy of 90.8%, 95.8%, and 80.3%, respectively) was higher than rate-based identification (overall accuracy of 27.5%, 15.0%, and 21.1%, respectively). Even when timing-based overall accuracy was relatively low (examples 2 and 5, 44.4% and 22.8%, respectively), it outperformed rate-based identification (31.7% and 10.3%). Whereas overall accuracy was a useful metric to describe the confusion matrices, it can be expected to be low relative to vowel-specific accuracies. Given twelve possible outcomes, an identifier operating at chance would have an overall accuracy of 8.3%. Thus, identification such as that based on the temporal responses of example neuron 3 (44.4%) were much greater than chance.

Systematic trends in overall accuracy versus RVF features were evaluated using principal component analysis of the RVFs. Principal components 1 through 3 (PC1, PC2, and PC3) were computed for the population of neurons for which both RVF and vowel responses were collected (Fig. 3, left). Together, PC1, PC2 and PC3 explained 95.0% of the RVF variance.

**Figure 3.**
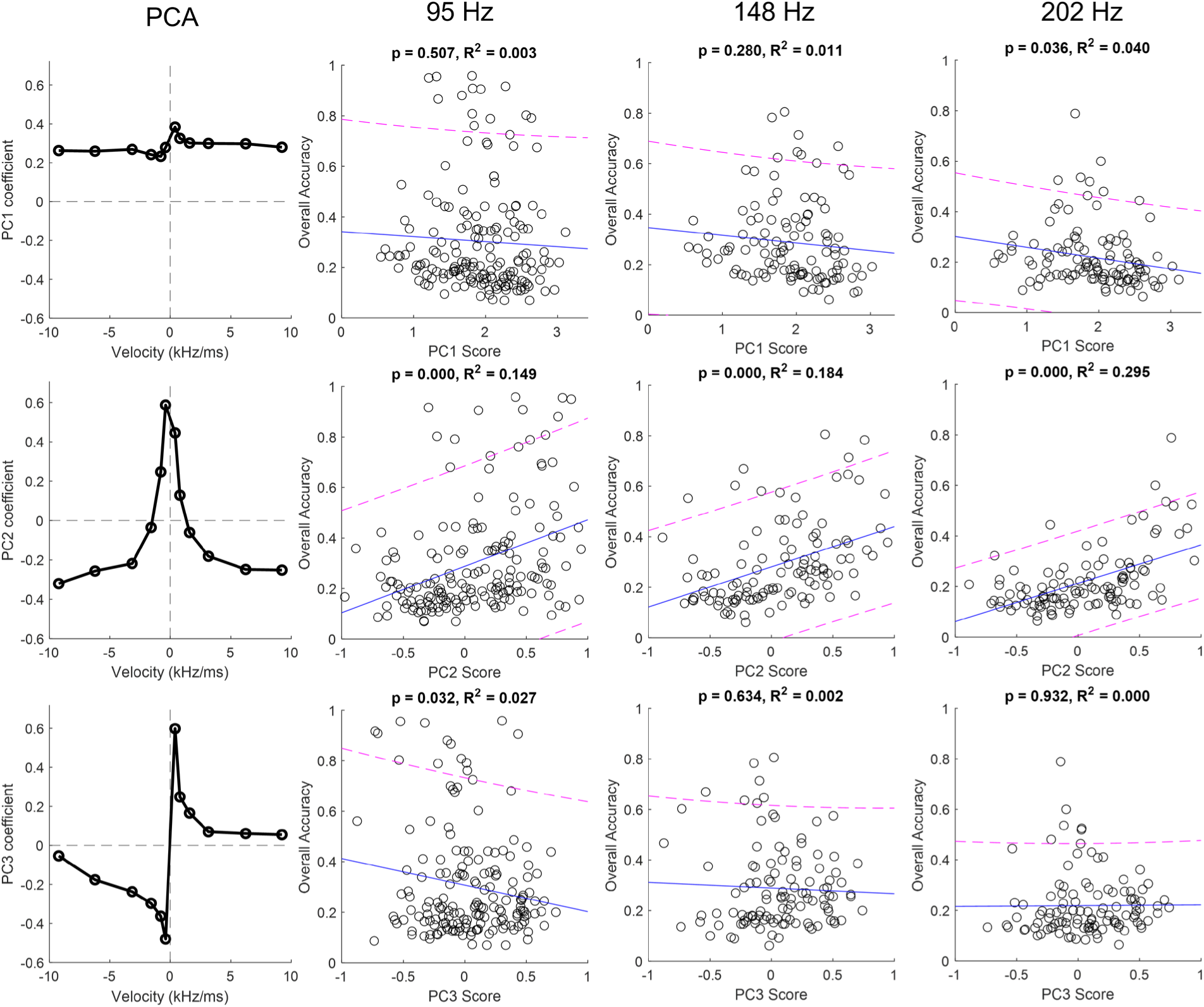
Principal component analysis (PCA) of neural RVFs. Scatter plots and regression analysis compare overall accuracy based on timing to principal-components score. Left column: Principal components 1, 2 and 3, plotted on loadings-versus-velocity axes. Dashed lines mark zero on both axes to assist PCA interpretation. Scatter plots show overall accuracy versus principal component scores for individual neurons, organized by speaker-F0 (columns) and principal component (rows). Blue regression lines reflect trends in data; magenta dashed lines are 95% confidence intervals. For each fit, p-values are from an ANOVA testing the hypothesis that the slope coefficient of the linear fit is non-zero. *RR*^2^ reflects variance explained by the linear fit.

PC1 (Fig. 3, left, top) reflected an overall average-rate feature, explaining 88.2% of the RVF variance, suggesting that the most prominent difference between RVFs was average rate. All loadings for PC1 had positive values, indicating that all rates varied together in PC1. PC2 (Fig. 3, left, middle) had a tendency for low-velocity rates to covary, and to vary inversely with high-velocity rates. In other words, neurons with high PC2 tended to exhibit high rates in response to low-velocity chirps and low rates in response to high-velocity chirps, regardless of chirp direction. PC2 explained 4.4% of RVF variance. Finally, PC3 (Fig. 3, left, bottom) reflected RVF direction-sensitivity, especially at low velocities. Neurons with high PC3 were upward-chirp biased. PC3 explained 2.5% of RVF variance. These results show that RVF rates varied based on absolute velocity, that is, velocity of chirps irrespective of direction, more than on chirp direction.

Quantifying principal component scores for each neuron revealed trends between RVF features and overall accuracy. Scatter plots showing overall accuracy versus principal component scores were separated by vowel F0s (columns in Fig. 3). For each, a regression line was fit to the data, and the quality of fit was described by a p-value (ANOVA, testing the null hypothesis that linear slope coefficient ≠ 0) and R^2^ (coefficient of determination, describing variance explained). Most notably, PC2 was correlated with overall accuracy for all speakers. The p-values were consistently < 0.001—furthermore, R^2^ indicate 14.9%, 18.4%, and 29.5% variance explained for vowel accuracies for vowels with average F0 equal to 95, 148, and 202 Hz, respectively. In comparison to PC2, PC1 and PC3 show little to no correlation with overall accuracy. However, note that PC1 vs. vowel accuracy for the 202-Hz F0 vowels and PC3 vs. vowel accuracy for the 95-Hz F0 vowels had p-values < 0.05. The overall accuracy values illustrated in Fig. 3 were for vowel identification based on neural timing. In comparison, there were no notable trends in overall accuracy based on average rate versus principal-component scores (not shown).

High chirp-direction bias at low chirp-velocities (PC3) was not significantly related to overall accuracy (Fig. 3, bottom). On the contrary, neurons for which low chirp-velocities elicited high response rates irrespective of chirp direction (i.e. neurons with high PC2 scores) tended to have higher overall accuracy. However, an alternative metric was necessary to directly assess the impact of high-velocity chirp direction bias on accuracy. The high-velocity direction bias metric (see Methods section 2.4) quantified the average direction bias in the RVF rates to the three fastest chirp-velocity pairs, ±3.16, ±6.24, and ±9.24 kHz/ms. Trends in overall accuracy versus high-velocity direction bias, based on both rate and timing, are illustrated in Fig. 4.

**Figure 4.**
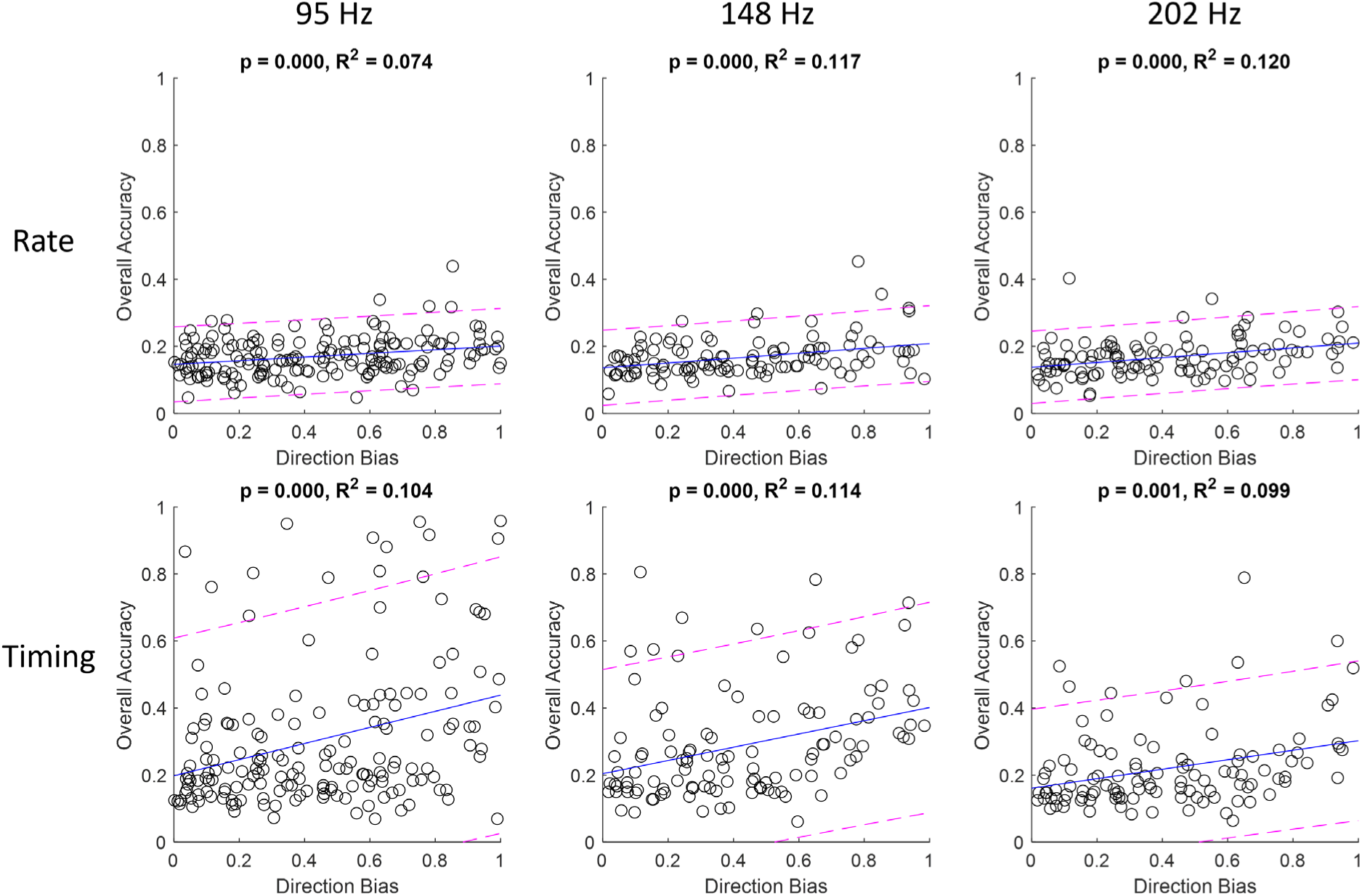
Scatter plots and regression analysis of overall accuracy (based on rate and timing identification) versus high-velocity direction bias. Plots are organized by speaker F0 (column) and neural metric (row). Blue regression lines reflect trends in data trend; magenta dashed lines are 95% confidence intervals. For each fit, p-values are the result of an ANOVA testing the hypothesis that the slope coefficient of the linear fit is non-zero. *RR*^2^ reflects variance explained by the linear fit.

There was a positive trend in high-velocity direction bias versus overall accuracy (p < 0.001 for all F0s; Fig. 4, top). However, the amount of variance explained by this metric was low, at 7.4%, 11.7%, and 12.0% for 95-Hz, 148-Hz, and 202-Hz F0s, respectively. Similarly, high-velocity direction bias was positively correlated with overall accuracy based on timing (Fig. 4, bottom; p < 0.01 for all F0s), but the percentage of variance explained by timing-based accuracy was also low, at 10.4%, 11.4%, and 9.9% for 95-Hz, 148-Hz, and 202-Hz F0s, respectively. However, note that rate-based accuracy was consistently low, whereas some individual neurons had near-100% overall accuracy based on timing.

Vowel identification was assessed for chirp-sensitive IC models (Mitchell and Carney, 2024) for comparison with the trends described above in the individual (Fig. 2) and population (Figs. 3-4) physiological data. Response profiles and confusion matrices are shown for three model IC neurons with 2-kHz CFs (Fig. 5), one biased towards upward chirps (top row), one biased towards downward chirps (middle row), and one without chirp direction bias (non-selective, NS) (bottom row). Model parameters are provided in Table I.

**Figure 5.**
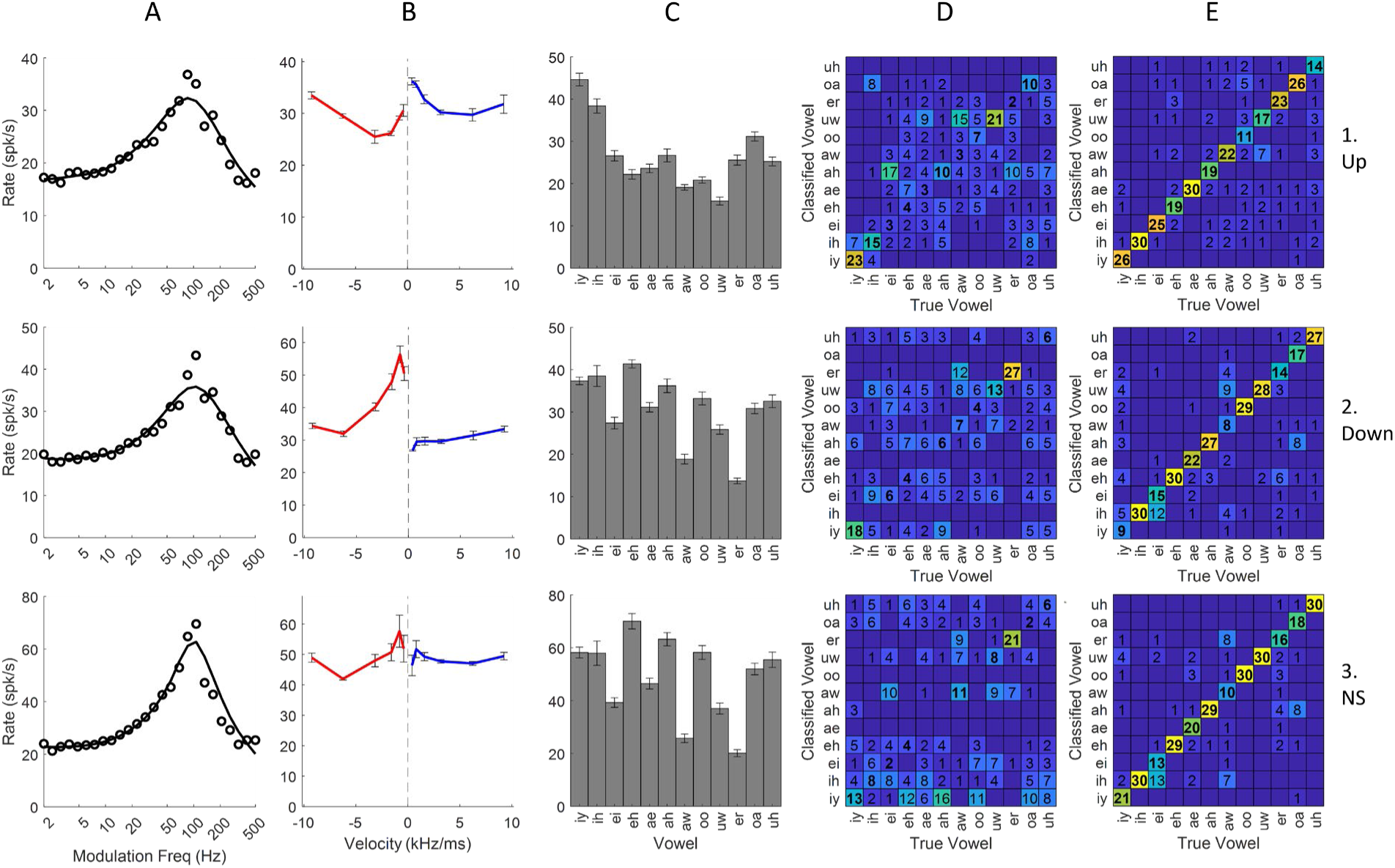
Response profiles of three model IC neurons, all with 2-kHz CFs but differing in chirp sensitivity due to octopus-cell inhibition. Top row) Upward-biased model neuron that received downward-biased octopus-cell inhibition. Middle row) Downward-biased model neuron that received upward-biased octopus-cell inhibition. Bottom row) Non-selective (NS) model neuron that received no octopus-cell inhibition. Stimulus responses are organized as in Fig. 2.

The chirp-selective model used here (Mitchell and Carney, 2024) is limited to neurons with BE MTFs and BMFs around 100-Hz (Fig. 5A). This BMF is near the F0 of vowel stimuli used to generate the confusion matrices shown, for vowels with F0 equal to 95 Hz. Simulated RVFs (Fig. 5B) differed solely in the presence or nature of inhibition from the octopus-cell model. The upward and downward biased neurons received inhibition from octopus-cell model cells that were downward and upward based, respectively. The non-selective neuron (bottom row) received no inhibition from the octopus-cell stage.

The average-rate profile across the set of vowels for the downward-biased model (Fig. 5C, Row 2) was similar to that of the non-selective neuron (Row 3). The upward-biased model rate profile (Row 1) had relatively larger differences from that of the non-selective model, with larger differences between the rate minima and maxima, among other dissimilarities. In general, average rates across the set of vowels differed across the models. In order to compare vowel identification between model neurons, and also to compare these model neurons to a similar physiological example (Fig. 2, Row 1), model rate functions were normalized to the average vowel rate of the physiological example (see Methods).

Average rate-based confusion matrices for the model neurons (Fig. 5D) were generated for responses to vowels with F0 equal to 95-Hz. Vowels with the greatest number of true positives tended to have larger extrema in the rate profiles, as expected. Timing-based confusion matrices (Fig. 5E) showed that specific vowel identifications differed in accuracy across the models. For identifications based on average rates, /er/ was identified reliably for the downward-biased and non-selective models, whereas the upward-biased model identified /iy/ and /uw/ more reliably. Accuracy of identification based on timing improved for some vowels, such as /aw/, in the upward-biased model neuron compared to the downward-biased or non-selective model neurons.

Overall accuracies for rate-based identification (28.1%, 25.3%, and 20.9% for upward, downward, and non-selective model neurons, respectively) were low compared to timing-based identification (72.8%, 71.1%, and 76.7% for upward, downward, and non-selective models, respectively) for these model neurons. In comparison to the non-selective model, the addition of octopus-cell inhibition benefited rate-based identification but slightly decreased the overall accuracy based on timing.

Response profiles and confusion matrices for three model IC neurons with 1-kHz CFs are shown in Fig. 6 (top row, upward-biased; middle row, downward-biased; bottom row, non-selective). Average rates for these models in response to vowels were normalized by that of Neuron 4 in Fig. 2.

**Figure 6.**
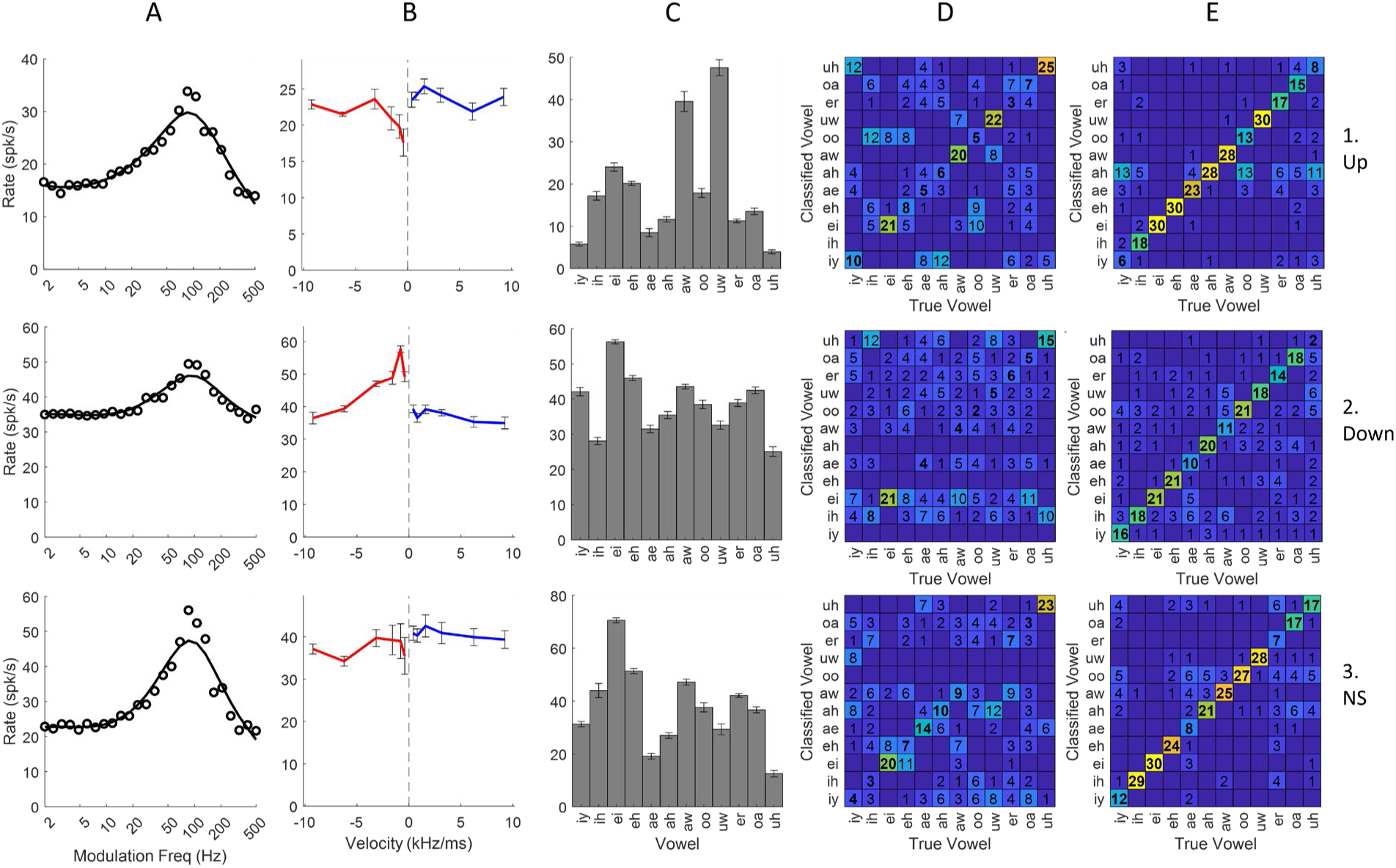
Response profiles of three model neurons, all with 1-kHz CF. Neurons differ by octopus-cell inhibition and RVF bias (upward, downward, and NS are shown on the top, middle, and bottom rows, respectively). Stimulus responses are otherwise organized identically to Figure 2.

The model MTFs were BE (Fig. 6A), and for these lower-CF (1-kHz) models, the direction bias in the model RVFs (Fig. 6B) was not as pronounced as for the 2-kHz model (Fig. 5), consistent with prior modeling results (Mitchell and Carney, 2024). Nonetheless, the three model neurons display markedly different rate profiles in response to vowels (Fig. 6C), despite their similar MTFs and identical CFs. In the rate-based confusion matrices (Fig. 6D), the rate profiles again were consistent with rate-based identification accuracy. Note that all model neurons had high identification accuracy for /ei/ and /uh/, but the upward-biased neuron also had high accuracy for /aw/ and /uw/. Timing-based confusion matrices (Fig. 6E) had vowel-by-vowel differences in accuracy as well; /ae/, /aw/ and /uh/ were particularly prominent examples. The overall accuracies for vowel identification varied across the models for rate-based (36.7%, 19.7%, and 30.5% for upward-, downward- and non-selective, respectively) and timing-based (70.0%, 54.1%, and 68.1%, for upward-, downward-, and non-selective, respectively. As was true for the 2-kHz models (Fig. 5), the upward-biased chirp-selective model outperformed the downward-biased model for both rate- and timing-based vowel identification; however, in the case of the 1-kHz models, addition of chirp selectivity had mixed effects on overall accuracy. Thus, taking both the 1-kHz and 2-kHz examples into account, adding chirp selectivity to the models had clear effects on the rate and timing responses of the models to the set of vowels, but there were not clear trends in the resulting changes in vowel-identification accuracy in these limited examples.

## 4. Discussion

We tested the hypothesis that chirp sensitivity of IC neurons is associated with discriminability of natural vowel tokens based on IC responses. Using a strategy of component decomposition, we identified within-pitch-period chirps in vowel stimuli with velocities in the range for which IC neurons have chirp selectivity (Mitchell et al., 2023). Confusion matrices generated based on rate and timing metrics showed that vowel identification based on a temporal analysis of IC responses was consistently more accurate than identification based on average rates. Vowel identification was not readily predicted by the RVF alone; however, a population-level analysis revealed a relationship of overall vowel-identification accuracy based on timing and Principal Component 2 (PC2), a chirp-directionless feature of RVFs related to a rate-bias toward low-velocity chirps in comparison to high-velocity chirps. No relationship between low-velocity chirp-direction bias (PC3) and overall vowel-identification accuracy was found; however, high-velocity chirp bias was significantly related to overall accuracy for vowel identification based on both rate and timing analyses.

The relationship between neural chirp selectivity and vowel responses could be further tested by collecting responses to vowels spoken with a larger set of F0s, possibly including multiple tokens of vowels spoken by each speaker, and then analyzing the ability to classify the vowels based on neural responses. This analysis would require a larger physiological dataset than was collected here.

Finally, a chirp-velocity sensitive computational IC model was used to examine the vowel-identification performance of model cells with and without octopus-cell inhibition. Rate profiles across the set of vowels differed between otherwise identical model neurons that received inhibition from octopus-cell models with different direction biases to introduced different chirp-velocity sensitivities in the IC model. Velocity sensitivity impacted model response profiles across natural-vowel tokens, though further work is needed to better match the entire set of model response features to specific examples of IC neurons studied physiologically.

Another method of interrogating the impact of chirp-sensitivity on vowel representation is to directly compare model neuron results. The 2-kHz CF models had vowel-rate histograms (Fig. 5C) that differed more between upward and non-sensitive neurons than between downward and non-sensitive neurons. This difference was corroborated by the rate-based confusion matrices (Fig. 5D), which featured different salient vowels, as well as by the timing-based confusion matrices (Fig. 6E), in which the upward-biased neuron displayed markedly different identification accuracies than the downward or non-sensitive models for the vowels /aw/, /oo/, and /ei/, among others. It is interesting to note that the vowel /aw/ (95-Hz F0) contains an upward chirp between 1.38 kHz and 2.81 kHz (Fig. 1); the presence of this cue potentially led to an elevated accuracy based on timing for the upward-biased, 2-kHz model.

A question this work attempted to answer is whether fast spectrotemporal chirps contained in vowels, which result from the phase differences between harmonics associated with resonances of the vocal tract, contributes to the representation of formant frequencies of natural vowels in the midbrain. Results of analysis on vowel responses of individual neurons in physiology alone are inconclusive, due to the inability to make controlled comparisons between IC neurons that differed only in chirp-direction sensitivity. Responses of model neurons allowed comparison of responses that differed in chirp sensitivity, and the model responses were compared to specific physiological examples. For example, the exemplar 2-kHz CF physiological neuron (neuron 1 in Fig. 2) had a BE MTF and a downward biased RVF, thus its vowel-rate histogram was expected to be similar to a model neuron with similar response properties. However, comparison of this unit’s vowel-rate profile to that of model neuron 2 in Fig. 5 shows that the rate profiles do not match. There was a similar lack of detailed correspondence between physiological neuron 4 (Fig. 2) and the 1-kHz model neuron 2 (Fig. 6), and between physiological neuron 5 (Fig. 2) and the 1-kHz model neuron 1 (Fig. 6). One would expect basic response properties of CF and MTF shape to drive vowel rate profiles, such as proximity of CF to formant frequencies. The fact that vowel responses of physiological and model neurons that share these properties differed, regardless of chirp-velocity sensitivity, suggest that there are additional factors unaccounted for in the IC model.

Of the two RVF features that were significantly correlated with overall vowel-identification accuracy, PC2 reflected a chirp-directionless property, describing RVF rates that are relatively high in response to low chirp velocities and low in response to high chirp velocities. While individual neurons with high PC2 scores may still be direction-biased for some velocities, this result suggests that absolute velocity (i.e. speed) of chirp was more indicative of vowel-identification accuracy than was direction bias. It is interesting to consider that the shape of PC2 would match the RVF of a simple hypothetical neuron that responded only to energy at CF: chirps that take a long time to pass through CF (i.e. low-velocity chirps) would elicit relatively higher rates than faster (high-velocity) chirps. In the aperiodic chirp stimulus with which RVFs are constructed, energy between chirps of different velocities was normalized; however, this normalization may not completely counteract this effect. However, the effect of high-velocity direction bias on overall vowel-identification accuracy (Fig. 4) should not be understated, suggesting that direction bias remains an important aspect of RVFs. A future version of the IC model should more precisely describe directionless chirp-velocity and high-velocity direction bias of neurons to better reflect these trends in the physiological results.

## Glossary – List of Abbreviations

AM: amplitude modulation
AN: auditory nerve
BE: band-enhanced
BMF: best modulation frequency
BS: band-suppressed
CD: coincidence detector
CF: characteristic frequency
DPOAEs: distortion product otoacoustic emissions
F0: fundamental frequency
F1: first formant
F2: second formant
IC: inferior colliculus
ICC: central nucleus of the inferior colliculus
MTF: modulation transfer function
NHPP: non-homogeneous Poisson process
OCF: off characteristic frequency
PCA: principal component analysis
PVCN: posteroventral cochlear nucleus
RIS: rate-independent spike (distance)
RM: response map
ROC: receiver-operating characteristic
RVF: rate-velocity function
VNLL: ventral nucleus of the lateral lemniscus

## Acknowledgements

We acknowledge the help of Kristina Abrams with physiological experiments, Douglas Schwarz with software, and Daniel Pyskaty with vow analysis. Constructive comments were provided by graduate committee members Professors Zhiyao Duan and Mark Bocko. Professor Ken Henry provided detailed comments on a pervious version of the manuscript. This work was supported by NIH R01-DC001641 and F31-DC019816.

